# Q-BioLiP: A Comprehensive Resource for Quaternary Structure-based Protein–ligand Interactions

**DOI:** 10.1101/2023.06.23.546351

**Authors:** Hong Wei, Wenkai Wang, Zhenling Peng, Jianyi Yang

## Abstract

Since its establishment in 2013, BioLiP has become one of the widely used resources for protein–ligand interactions. Nevertheless, several known issues occurred with it over the past decade. For example, the protein–ligand interactions are represented in the form of single chain-based tertiary structures, which may be inappropriate as many interactions involve multiple protein chains (known as quaternary structures). We sought to address these issues, resulting in Q-BioLiP, a comprehensive resource for quaternary structure-based protein–ligand interactions. The major features of Q-BioLiP include: (1) representing protein structures in the form of quaternary structures rather than single chain-based tertiary structures; (2) pairing DNA/RNA chains properly rather than separation; (3) providing both experimental and predicted binding affinities; (4) retaining both biologically relevant and irrelevant interactions to alleviate the problem of the wrong justification of ligands’ biological relevance; and (5) developing a new quaternary structure-based algorithm for the modelling of protein–ligand complex structure. With these new features, Q-BioLiP is expected to be a valuable resource for studying biomolecule interactions, including protein–small molecule, protein–peptide, protein–protein, and protein–DNA/RNA. Q-BioLiP is freely available at https://yanglab.qd.sdu.edu.cn/Q-BioLiP/.

## Introduction

The biological functions of many proteins are achieved by interacting with other biomolecules, which are referred to as ligands. A collection of high-quality data for protein–ligand interactions is essential to enable related computational studies [1–5], such as in the prediction of the protein–ligand binding site [6–11], prediction of binding affinity [12, 13], and protein–ligand docking [14–16]. BioLiP is a database that collects 3D structures of biologically relevant protein–ligand interactions [17]. It has emerged as one of the most widely used resources for investigating protein–ligand interactions.

There are several known inherent issues with the data in BioLiP. The first one is that the protein structures are represented in single chain-based tertiary structures. Nonetheless, the functional form of many proteins is in quaternary structures, typically comprising multiple interacting chains. Consequently, some vital protein–ligand interactions are not captured in the BioLiP data due to the incompleteness of the protein structure [18], particularly when a ligand simultaneously interacts with multiple protein chains. For example, the hemoglobin protein can only transport oxygen in the form of a tetramer, which is composed of four chains. Second, DNA ligands in BioLiP are presented in a single-chain format, which is inconsistent with the fact that DNAs typically form double-helix structures consisting of two complementary chains. The third one is the potential misjudgement of the biological relevance. BioLiP employs an empirical rule to determine whether a protein–ligand interaction is biologically relevant or not. Consequently, a protein–ligand interaction that is deemed biologically irrelevant is excluded from BioLiP, resulting in the problem of missing data, when a biologically relevant interaction is mistakenly judged as irrelevant.

In this article, we introduce an enhanced version of BioLiP, named Q-BioLiP, to address the aforementioned issues. The major updates include the representation of protein structures in quaternary structures, the retention of both biologically relevant and irrelevant interactions, the paired chain structures for DNA/RNA ligands, the adoption of Macromolecular Crystallographic Information File (mmCIF) format [19] rather than Protein Data Bank (PDB) format [20] to deal with huge molecules (that exceed the capability of the PDB format), and the inclusion of computed binding affinity. Last but not least, we provide an efficient template-based approach to ligand-binding site prediction using the Q-BioLiP data, which allows for the structure input consisting of either single chain or multiple chains.

## Database Construction

### Overview of Q-BioLiP

The flowchart for building Q-BioLiP is presented in **Figure 1**. First, starting from the asymmetric unit files (in mmCIF format), quaternary structures (known as biological units) are generated using the rotation and translation matrices stored in the mmCIF files. Then, receptors and ligands are extracted from the quaternary structures. In this work, receptors are defined as proteins consisting of ≥ 30 amino acid residues (AAs) in each polypeptide chain, while other molecules are defined as ligands (including small molecules, peptides consisting of < 30 AAs, and DNA/RNA). As base pairing can be formed between DNA/RNA chains, a heuristic algorithm (see “An effective DNA/RNA pairing algorithm” in this section) is proposed to pair the DNA/RNA chains. As done in BioLiP [17], the biological relevance of each ligand–protein complex is assessed with a semi-manual procedure. For the sake of completeness, Q-BioLiP also retains quaternary structures with ligands or receptors only. Thus, the data deposited in Q-BioLiP fall into three categories: biologically relevant ligand–protein interactions, biologically irrelevant ligand–protein interactions, and structures with ligands or proteins only. They can be accessed through a user-friendly web interface at our website: https://yanglab.qd.sdu.edu.cn/Q-BioLiP/.

**Figure 1.**
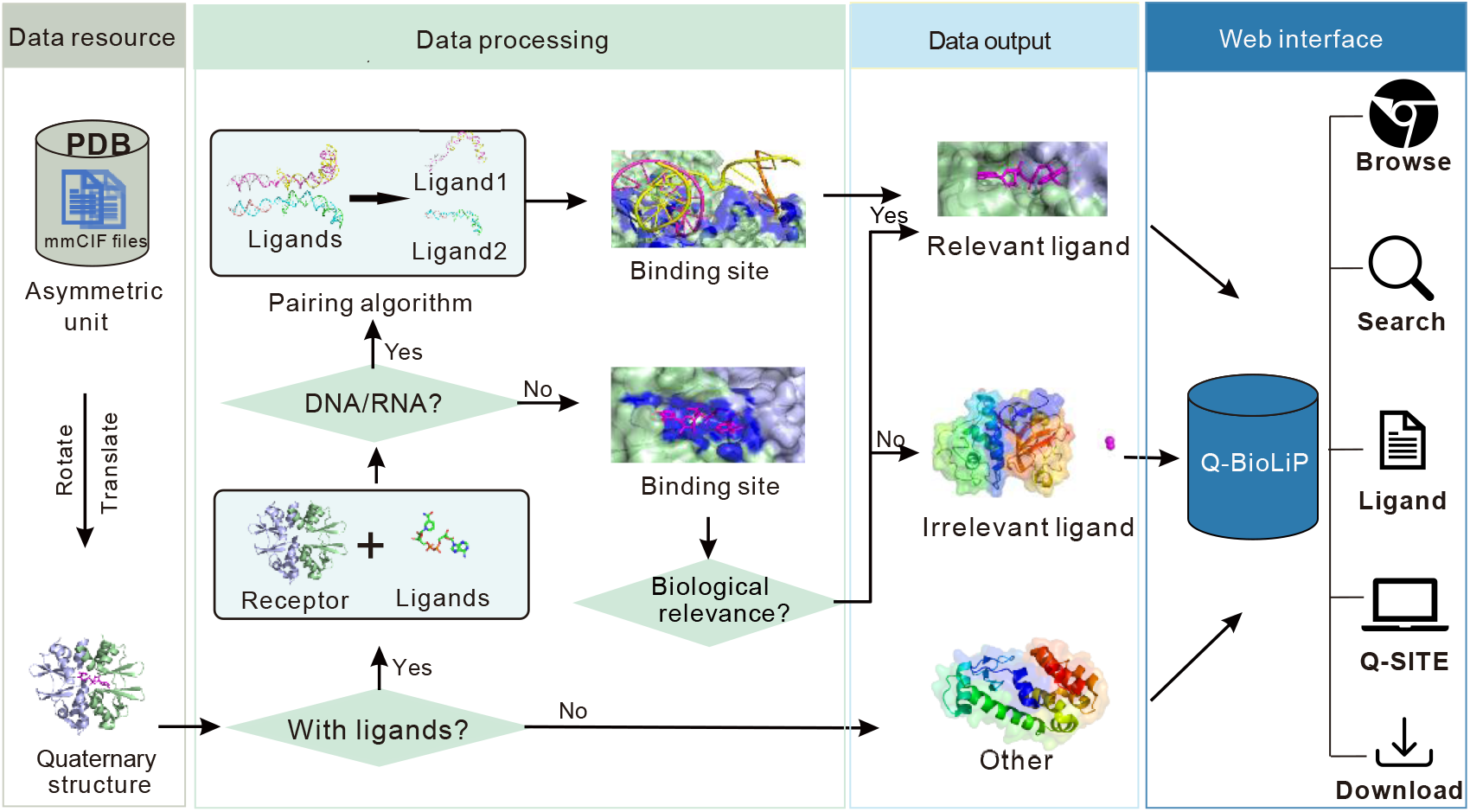
Workflow of Q-BioLiP. Quaternary structures are first generated from the asymmetric unit files in PDB. The quaternary structures are then processed into three categories, biologically relevant interactions, biologically irrelevant interactions, and structures with ligands or proteins only. All the data are accessible through a web interface. PDB, Protein Data Bank.

### Procedure for database construction

The raw data are downloaded from the PDB database [20]. The procedure for the construction of Q-BioLiP consists of three major steps.

#### Generation of quaternary structures

For each PDB entry, the asymmetric unit (in mmCIF format) is first downloaded. One or more quaternary structures are generated according to the rotation matrices and the translation vectors stored in the mmCIF file’s records, *i.e.*, ‘pdbx_struct_assembly’, ‘pdbx_struct_assembly_gen’, and ‘pdbx_struct_oper_list’. Modified residues are converted to standard ones based on the record ‘pdbx_struct_mod_residue’. For Nuclear Magnetic Resonance (NMR) structures, only the first model is taken into consideration in the procedure.

#### Extraction of receptor and ligands from quaternary structures

Each quaternary structure is separated into one receptor and multiple ligands (when they exist). The receptor consists of protein chains with ≥ 30 AAs; while ligands can be small molecules, DNA/RNA, or peptides with < 30 AAs.

#### Annotations of protein–ligand interactions

The interaction between the receptor and each ligand is annotated in terms of binding residues, biological relevance, binding affinity, and area of binding surface.

##### Binding residues

Binding residues are determined based on the atomic distances between the protein and ligand atoms. A residue is considered a binding residue if its closest atomic distance to the ligand is less than a defined threshold, which is the sum of the Van der Waal’s radius of the corresponding atoms plus 0.5 Å [21].

##### Biological relevance

Small molecules that are utilized to help determine the structure of proteins but do not have any biological activity are considered biologically irrelevant. Peptides and nucleic acids are regarded as biologically relevant to receptors without the need for assessment. We adopt a similar semi-manual procedure used in BioLiP to assess the biological relevance of the small molecules. Briefly, a list of 465 small molecules that are commonly used in protein structure determination is collected manually (available at the *Download* page). Small molecules outside this list are regarded as biologically relevant by default. For molecules in this list, a hierarchical procedure is employed, which involves computing of the number of binding residues, text mining of the PubMed abstract, and manual verification of suspicious entries. As a result of utilizing quaternary structure in this context, two of the criteria that are used in BioLiP, namely the ligand occurrence number and the continuity of binding residues, are deactivated. More details can be found in the BioLiP paper.

##### Binding affinity

The strength of ligand–protein interaction is commonly referred to as binding affinity. Experimental ligand-binding affinity is collected from three databases: Binding MOAD [2], PDBbind [3], and BindingDB [4]. However, only a limited portion (9.8%) of PDB entries have experimental binding affinity data. To address this issue, predicted binding affinity data are provided in Q-BioLiP for entries that do not have experimental binding affinity data. The predicted binding affinity data are obtained based on a few well-established physics-based tools, including X-Score [22], ITScore [14], and AutoDock Vina [16]. To generate a consensus binding affinity prediction, a regression model was trained to fit the real binding affinity by taking the three scores as input.

##### Area of binding surface

Besides binding affinity, the area of the binding interface is provided in Q-BioLiP. Motivated by Dockground [23], we measured the binding interface area *S* as below:

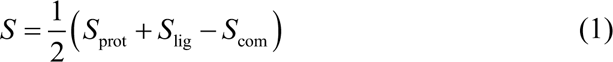

where *S*_prot_, *S*_lig_, *S*_com_ are the solvent-accessible surface area calculated by the program FreeSASA [24], with the protein, ligand, and ligand–protein complex structures as inputs, respectively.

### An effective DNA/RNA pairing algorithm

As is well known, DNA typically forms double-helix structure with two complementary chains. Inter-chain interactions are also witnessed between different DNA and RNA chains. However, in BioLiP, the DNA/RNAs are simply divided into separated chains, making the DNA/RNA structures incomplete. Thus, it is necessary to correctly pair DNA/RNA chains to keep them intact.

To address the above issue, we propose a simple yet effective pairing algorithm for DNA/RNA chains. To pair the DNA/RNA chains, we first parse the secondary structure of the whole DNA/RNA structure using the DSSR program [25]. Then the pairing state of each nucleotide is derived from the output of DSSR. Two chains are paired together if there are at least three inter-chain base pairs. Furthermore, in the case of a chain consisting of one or two nucleotides, the pairing is preserved as long as all of its nucleotides are paired with another chain.

We have validated the algorithm using a randomly selected set of 100 structures (list available at https://yanglab.qd.sdu.edu.cn/Q-BioLiP/DATA/pair_100.txt). The ground truth was obtained through manual verification. The algorithm described above produces pairings that are identical to the ground truth, indicating its effectiveness. Nevertheless, incorrect pairings may also occur for RNAs with new pairing patterns. To deal with such issues, we will keep improving our algorithm.

## Data content and discussion

### Comparison with BioLiP

The major differences between the data in Q-BioLiP and BioLiP are summarized in

**Table 1**, including the following five aspects.

**Table 1.**
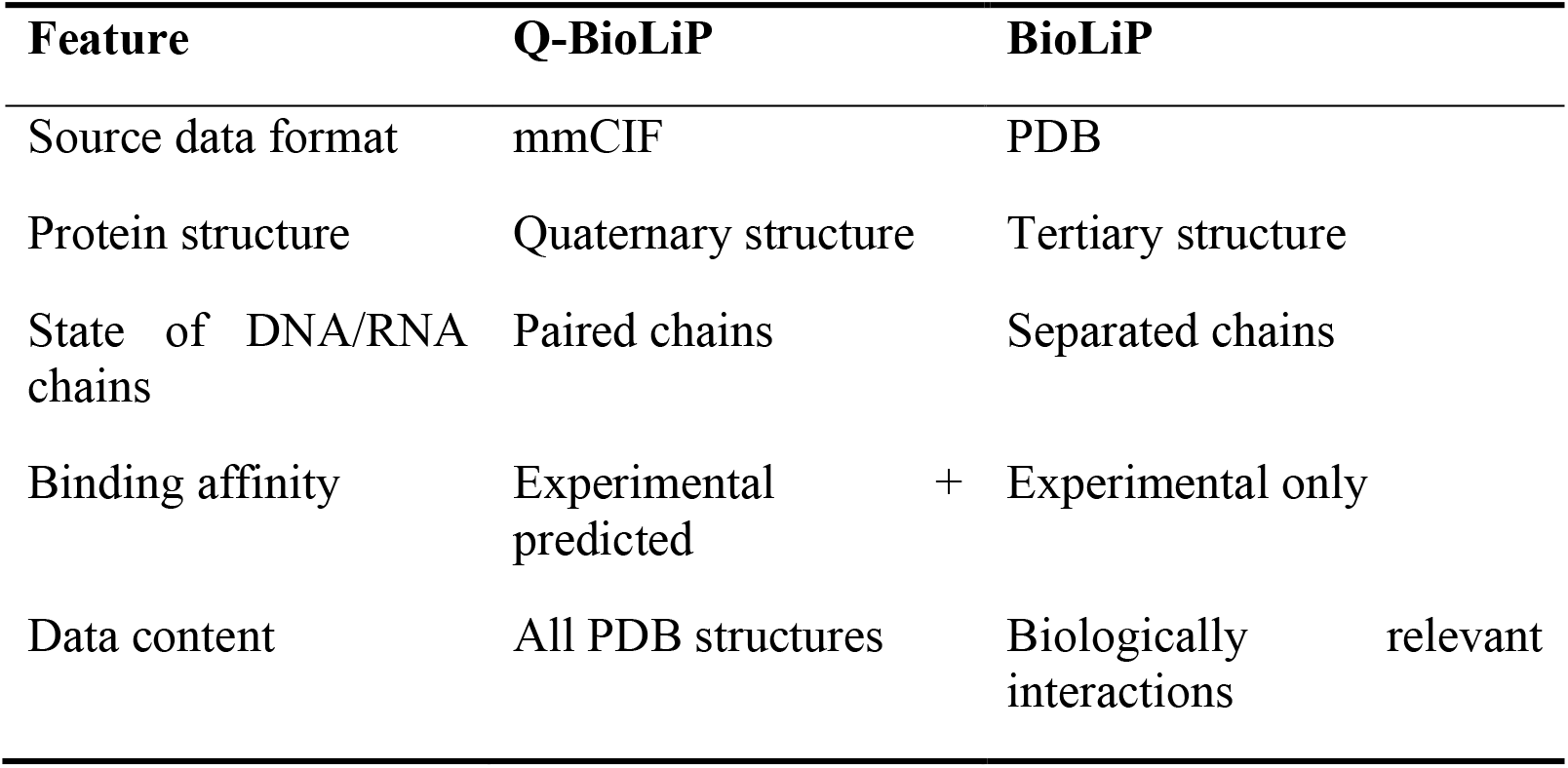
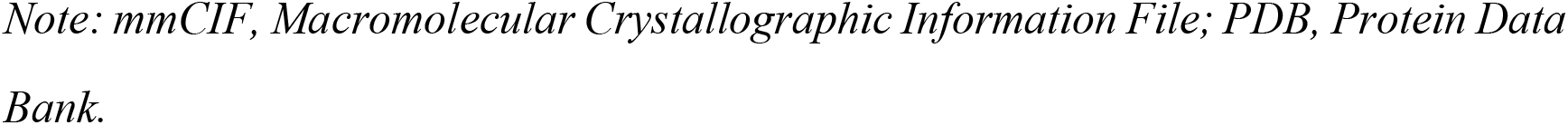
The major difference between the data in Q-BioLiP and BioLiP.

#### Source data format

Due to the limitation of the PDB format, structures with more than 62 chains such as many Cryo-Electron Microscopy (Cryo-EM) structures are ignored in BioLiP. The source data processed by Q-BioLiP is in mmCIF rather than PDB format, which solves the issue of missing structures in BioLiP.

#### Protein structure

One of the key developments of Q-BioLiP over BioLiP is the change in the form of protein structure. Q-BioLiP improves the representation of protein structure from single-chain based tertiary structure to quaternary structure. As indicated above, the quaternary structure may consist of single chain or multiple chains. The completeness of the protein structure can ensure the completeness of ligand-binding interactions [18]. For instance, **Figure 2A** shows the interaction between the HIV-1 protease and its inhibitor (PDB ID: 1EBY), where the protease is a C2 symmetric homo dimer and the catalytic residues are located on the interface. This interaction is however separated into two individual entries in BioLiP (each for one receptor chain, Figure S1).

**Figure 2.**
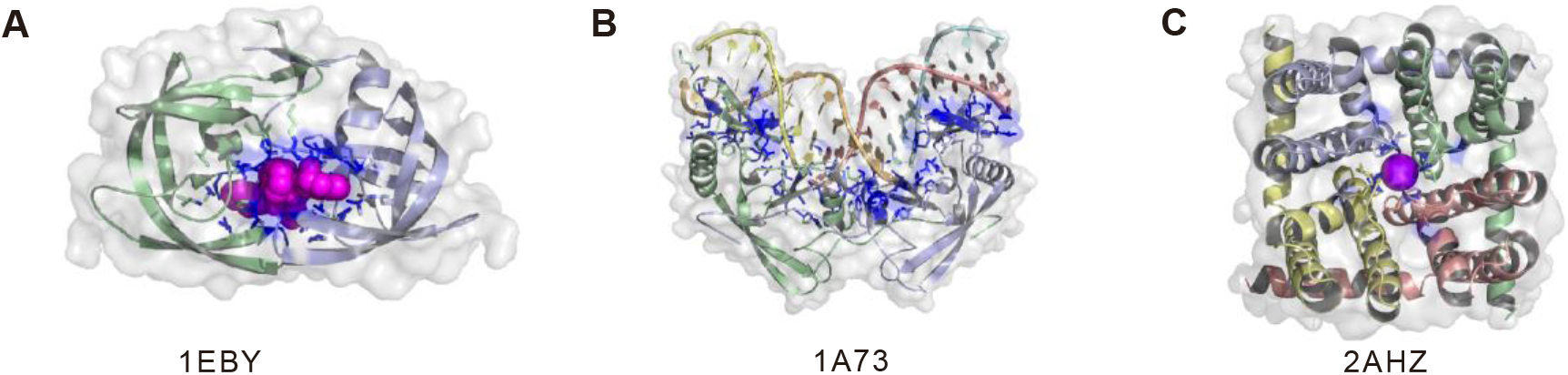
Three examples of protein–ligand interactions in Q-BioLiP. **A.** The interaction between the HIV-1 protease and its inhibitor (PDB: 1EBY). **B.** The DNA ligand binding with the structure of an intron encoded endonuclease (PDB: 1A73). **C.** The structure of the tetrameric NaK channel bound with K^+^ (PDB: 2AHZ).

#### State of DNA/RNA chains

With the pairing algorithm introduced above, we are able to correctly pair DNA/RNA chains. For example, the DNA ligand binding with the structure of an *intron encoded endonuclease* is formed by four DNA chains (PDB ID: 1A73, Figure 2B). On the contrary, BioLiP separates the interactions into 6 individual entries (see Figure S2): 2 receptor chains, and 3 DNA chains binding with each receptor chain (one DNA chain is ignored as it does not interact with the receptor chain after separation).

#### Binding affinity

In Q-BioLiP, ∼49,000 entries are annotated with experimental binding affinity data. In addition, predicted binding affinity and binding interface area are also provided for ∼1.7 million protein–small molecule interactions (peptide and DNA/RNA ligands are excluded).

#### Data content

Only interactions judged as biologically relevant are kept in BioLiP. However, this may result in the omission of some biologically relevant interactions due to misjudgement. For example, the structure of the *tetrameric NaK channel binding with K+* (PDB ID: 2AHZ) was judged as biologically irrelevant in BioLiP. In fact, the ion K+ is located in the pocket of a tetrameric cation channel and thus biologically relevant. This is correctly captured in Q-BioLiP due to the utilization of quaternary structure (Figure 2C). Around 64,000 Q-BioLiP entries (involving 12,000 PDB entries), initially filtered out by BioLiP, have been retained in Q-BioLiP due to the consideration of quaternary structure, and rule adjustments.

We admit that it is also possible that the judgement of biological relevance in Q-BioLiP is not perfect. We thus keep all data in our database so that the users can decide the relevance based on their own expertise. For the sake of completeness, structures with ligands or proteins only are also stored in our database, which may be used for other purposes, such as in protein structure database searching. To summarize, the Q-BioLiP data are organized into three categories, biologically relevant interactions; biologically irrelevant interactions; and structures with ligands or proteins only.

### Statistics of the Q-BioLiP data

By the time of October 11th, 2023, Q-BioLiP contains about 2.7 million entries, involving about 210,000 structures from PDB. These structures are determined by the following experimental methods (see Figure S3): X-radiation (X-ray, 85%), NMR (6.7%), Cryo-EM (8.2%), and others (1%). The overall distribution of the Q-BioLiP entries are shown in **Figure 3A**. Among the total entries, ∼35/63% are annotated as biologically relevant/irrelevant interactions (the orange/blue region); the remaining 2% are structures with ligands or proteins only (the black region). Further analysis is conducted on the biologically relevant interactions, which are the core data of Q-BioLiP.

**Figure 3.**
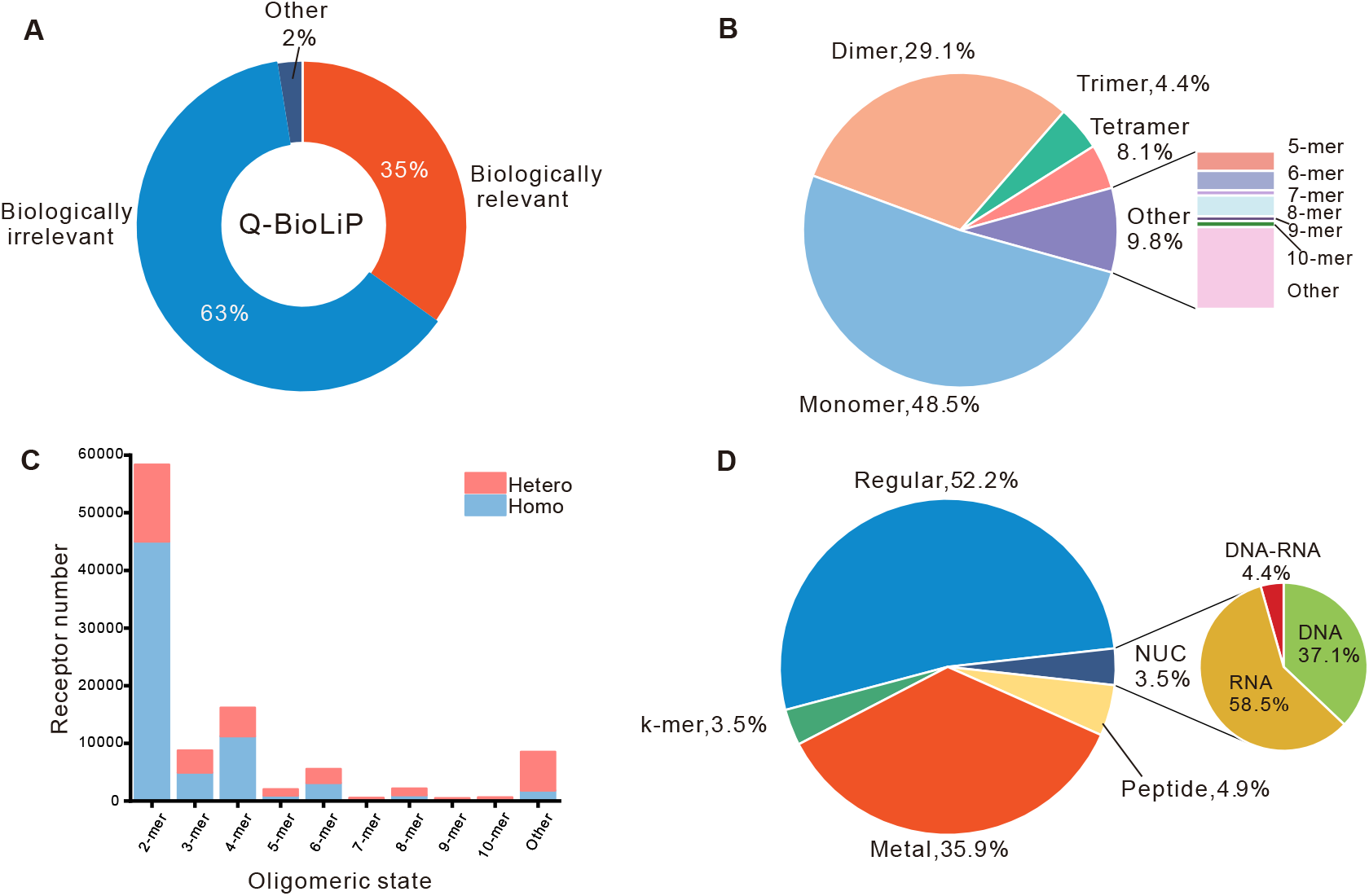
Statistics of the Q-BioLiP data. **A.** Overall statistics for the Q-BioLiP entries. **B.** Distribution of receptors in terms of the oligomeric state. **C.** Distribution of receptors in terms of homo- and hetero-oligomers at different oligomeric states. **D.** Distribution of ligands. Note that the data shown in panels B–D only involve the biologically relevant interactions. NUC, denotes the nucleic acids ligand.

The distribution of the proteins involved in biologically relevant interactions is shown in Figure 3B and C. Figure 3B suggests that almost half (48.5%) of these proteins are monomers, while the remaining half contain more than one chains (called oligomers). Interestingly, oligomers with even numbers of chains (*e.g.*, dimmer and tetramer) are more than those with odd numbers of chains (*e.g.*, trimer). The oligomers can be divided into two groups, homo-oligomers and hetero-oligomers, depending on if all chains are identical or not. Figure 3C shows the proportion of homo-oligomers and hetero-oligomers at different oligomeric states. For oligomers with even numbers of chains (especially for dimers and tetramer), the numbers of homo-oligomers tend to be higher than the numbers of hetero-oligomers.

Figure 3D shows the proportion of different ligand types involved in biologically relevant interactions. Regular small molecules account for more than half (52.2%) of all ligands; metal ions are the second largest group (35.9%). For nucleic acids ligands (3.5%), 58.5% and 37.1% of them are RNA and DNA, respectively; the remaining 4.4% are DNA–RNA complex.

Among the biologically relevant interactions involving oligomeric targets, 28.8% of the ligands interact with more than one chains (Figure S4A), which spreads across states from dimmers to 10-mers (Figure S4B). This indicates the necessity of updating BioLiP to Q-BioLiP, to maintain the completeness of ligand–protein interactions.

### Derivation of interaction-based data from Q-BioLiP

Besides protein–small molecule interaction, a variety of interaction data are derived from Q-BioLiP, which are shown in **Table 2**. These data can be easily used to construct training and/or benchmark datasets in the development of methods for protein–metal ion interaction [26], protein–peptide interaction [27], protein–protein interaction [28], protein–DNA/RNA interaction [29], and RNA–small molecule interaction [30]. These data are available for download at the Download page.

**Table 2.** Interaction-based data derived from Q-BioLiP.

| **Interaction type** | **Number of redundant interactions** | **Number of non-redundant interactions** |
| --- | --- | --- |
| protein–small molecule | ~460,000 (~140,000) | ~110,000 (~28,000) |
| protein–metal ion | ~300,000 (~79,000) | ~73,000 (~19,000) |
| protein–peptide | ~31,000 (~20,000) | ~8,000 (~4,000) |
| protein–protein<sup>*</sup> | ~150,000 (N/A) | ~38,000 (N/A) |
| protein–DNA | ~10,000 (~8,000) | ~3,000 (~2,000) |
| protein–RNA | ~12,000 (~6,000) | ~4,000 (~2,000) |
| RNA–small molecule | ~21,000 (~3,000) | ~4,000 (~200) |

*Note:* () denotes the number of receptors, <sup>\*</sup> refers to oligomer structures.

### Binding affinity data

About 49,000 entries in Q-BioLiP (from ∼21,000 unique PDB entries) are annotated with experimental binding affinity. Predicted binding affinities are provided for other entries without experimental binding affinity. Benchmark test on 285 high-quality data from the PDBbind core set (v2016) shows that the binding affinity predicted by these tools correlates well with the experimental binding affinity, with Pearson’s correlation coefficient (PCC) around 0.6 (Figure S5A–C). The consensus binding affinity prediction based on a regression model yields a 4.7∼15.5% higher PCC than the individual predictions (Figure S5D). Further tests based on a bootstrap sampling on the dataset by 1,000 times shows the improvement is statistically significant (Figure S6).

## Database interface

The Q-BioLiP database is freely accessible at https://yanglab.qd.sdu.edu.cn/Q-BioLiP/. Five modules are provided to use the data (illustrated in Figure 4), which are introduced below.

**Figure 4.**
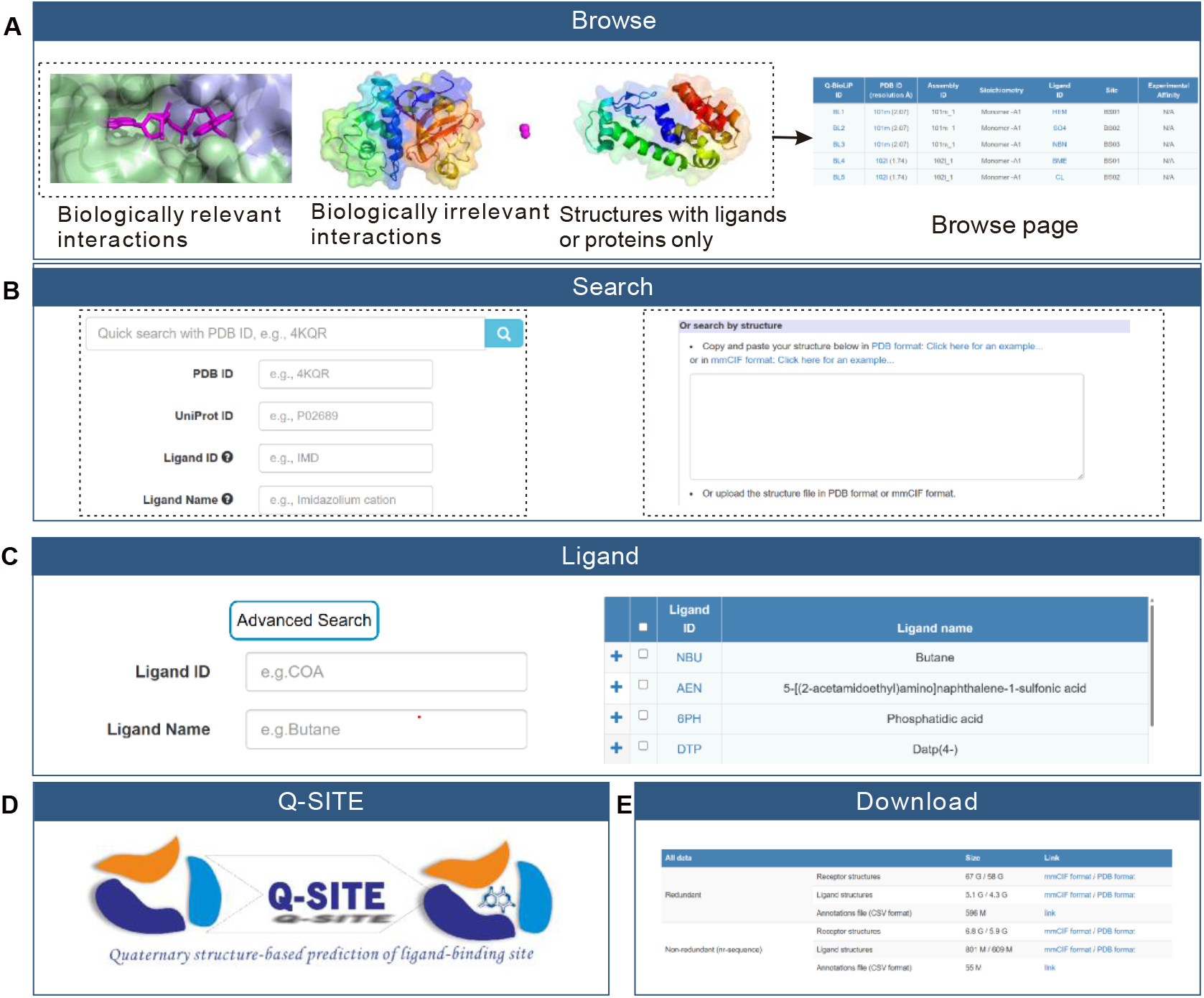
Major modules of the Q-BioLiP web interface. **A.** Through the *Browse* module, users can browse entries of three categories. **B.** Through the *Search* module, users can perform search with two options. **C.** Through the *Ligand* module, users can search for detailed information for the specific ligand or browse all ligands. **D.** A quaternary structure-based protein–ligand complex modelling module *Q-SITE* is provided, which supports structure inputs with both monomers and oligomers. **E.** Through the *Download* page, all data in Q-BioLiP can be downloaded freely.

### Browse module

There are three options in the browse module: (i) Browse biologically relevant entries, (ii) Browse biologically irrelevant entries, and (iii) Browse structures with ligands or proteins only. Clicking on each option would display the summary of entries in the corresponding category in the form of a table, which has seven columns: Q-BioLiP ID, PDB ID with resolution, Assembly ID, Stoichiometry, Ligand ID, Site and Binding affinity. The detailed information for each entry is available by clicking on the corresponding Q-BioLiP ID.

### Search module

In the search module, users can query the Q-BioLiP data using two options. The first one is a rapid search of the following columns: PDB ID, UniProt ID, Ligand ID, and Ligand Name. An input box is provided to filter results further. The results can be downloaded in several formats including JSON, CSV, and so on. Another one is protein structure-based search using the Foldseek algorithm [31], which usually takes ∼2 minutes to complete.

### Ligand module

In this module, all ligands involved in Q-BioLiP are displayed in two columns by default: Ligand ID and Ligand Name. More detailed information can be shown by clicking on the Ligand ID link. The detailed ligand page contains the 2D view of ligand, synonyms, as well as the chemical component summary. To sufficiently annotate ligands, the synonyms for each ligand are collected from PDB [20], ChEBI [32], and PubChem [33], which are also essential in the biological relevance assessment. The chemical component summary includes molecular weight, chemical formula, the SMILES string, and external links. Besides, a search module (the “Advanced Search” button) is provided for searching for a specific ligand by ligand ID or ligand Name.

### Quaternary structures-based binding site prediction module Q-SITE

Q-SITE is an extension of the COACH-D algorithm [6] for template-based modelling of protein–ligand complex structures. Q-SITE supports the structure inputs with both oligomers and monomers. The modelling is based on quaternary structure templates in Q-BioLiP, which is about 40 times faster and 6.5% more accurate than COACH-D. More details about Q-SITE will be introduced elsewhere. Users can provide either experimental structure or predicted structure as inputs, the server will return the top five predictions within ∼20 minutes on average.

### Download module

The Q-BioLiP data are provided for download at this module. Three tables are provided on the Download page: All data, Interaction-based data, and Weekly Update. In the ‘All data’ table, both redundant and non-redundant versions are provided for easy download. The redundant version contains ∼2.6 million entries of interactions (i.e., biologically relevant + irrelevant interactions) and ∼68,000 entries with proteins or ligands only.

For the core data of the biologically relevant interactions, two non-redundant datasets (denoted by nr-sequence and nr-structure) are created based on sequence and structure similarity, respectively. As quaternary structure may contain multiple chains, we define redundancy as follows. First, the protein structures are grouped by the oligomeric state (*i.e.,* the number of chains). Second, the chain-level sequences and structures are clustered by CD-HIT [34] and Foldseek [31], respectively. To identify redundant structures within each group, we compare pairs of chains from the two structures. If any pair of chains exhibits a similarity (*i.e.,* sequence identity or Foldseek score) higher than 0.9, those structures are considered redundant. The nr-sequence dataset has ∼41,000 receptors and ∼240,000 ligands; which are compared to ∼29,000 receptors and ∼160,000 ligands, respectively, in the nr-structure dataset. This suggests that protein structures are more conserved than protein sequences. Analysis shows that the distributions of the ligands binding to multiple chains or single chain are similar to the redundant dataset (Figure S4).

Within the ‘Interaction-based data’ table, users can download the previously mentioned interaction-based datasets. Given that some users may be unfamiliar with the mmCIF format, we provide a script to convert between mmCIF format and PDB format. This script is designed to automatically detect the input format and convert it to the other format. Q-BioLiP is updated weekly with PDB. The updated data are provided for download in the ‘Weekly Update’ table, to avoid re-download of the previous data. For users who are interested in monomer structure-based protein–ligand interaction, we also provide the redundant data in single-chain format like BioLiP. They are obtained by splitting the quaternary structures into individual receptor chains. The data are available for download at a separate link (https://yanglab.qd.sdu.edu.cn/Q-BioLiP/Download/index_biolip.html).

### Detailed information on protein–ligand interactions

The information on protein–ligand interactions is presented as a webpage, which consists of three tabs. The first tab is the 3D visualization of the interaction (Figure 5A). The visualization is powered by the 3Dmol package[35]. Two buttons are provided to show the local view and specific binding residues, respectively. In addition, the specific binding residues can be shown in a table by clicking on the “show binding residues” button. On the right of the page (Figure 5B), the overall information is organized into three sections: *Receptor*, *Ligand*, and *Download*. In the *Receptor* section, the stoichiometry type is derived with the same definition as in PDB, *i.e.*, two chains are considered equivalent if their sequence identity is greater than 95%. In the *Ligand* section, three items are shown: ligand ID, ligand name, as well as the biological relevance. In the *Download* section, the structures of receptor and ligand are provided in both mmCIF and PDB formats. Structures comprising more than 62 chains are currently unavailable in the legacy PDB format. In order to maintain completeness, these large structures are divided into multiple PDB files. A tarball (.tar.gz) is provided on the page, which is a bundle of these files along with a text file to map the original chain IDs to the new IDs.

**Figure 5.**
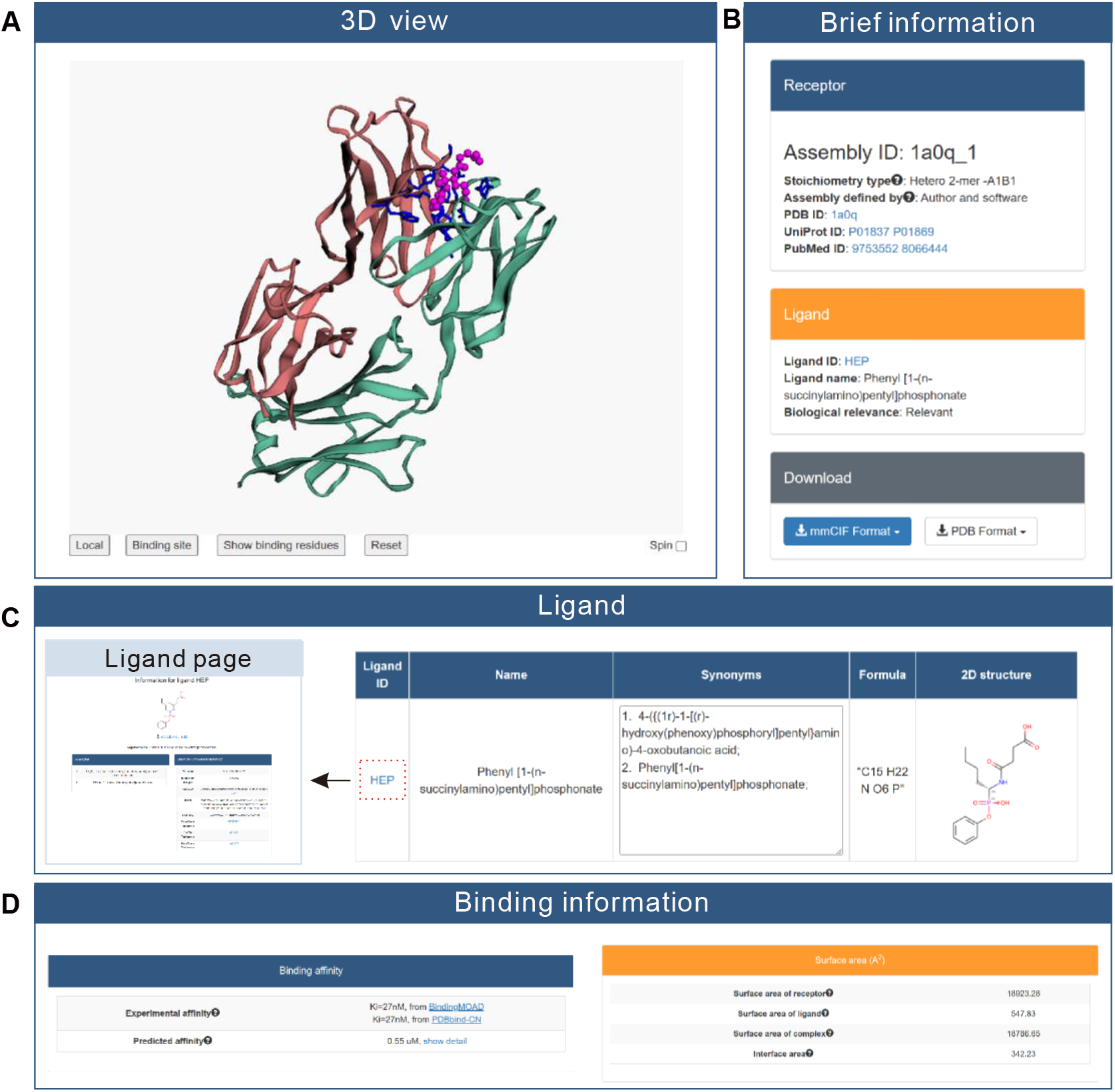
Information displayed on the webpage for an example Q-BioLiP entry. **A.** The 3D visualization of the ligand–protein interaction. **B.** The overall brief information about Receptor, Ligand, and Download. **C.** The detailed ligand information, including Ligand ID, Ligand Name, Synonyms, Formula, and 2D structure. **D.** The binding affinity and surface area.

The second tab is the ligand information (Figure 5C), including ligand ID (with a link to the detailed information page), ligand name, synonyms, formula, and 2D image of the ligand structure.

The last tab is the binding information (Figure 5D), which consists of the binding affinity and the surface area. Both experimental (if available) and predicted binding affinities are provided.

## Conclusion

We have developed the Q-BioLiP database for quaternary structure-based protein–ligand interactions. The major contributions include: (1) All structures in Q-BioLiP are based on quaternary structures, making the protein–ligand interactions complete; (2) DNA/RNA chains are properly paired; (3) Both experimental and predicted binding affinities are provided; (4) To address the problem of misjudgement of biological relevance, both relevant and irrelevant entries are kept; (5) A quaternary structure-based protein–ligand complex modelling algorithm is developed. We believe that the development of Q-BioLiP will better serve the community of protein–ligand interactions.

During the submission of this work at the beginning of July 2023, the authors noted an updated version of the original BioLiP (i.e., BioLiP2 [36], https://zhanggroup.org/BioLiP). Q-BioLiP and BioLiP2 were developed independently based on the original BioLiP database. They are complementary to each other in terms of both web interface and underlying data. Our Q-BioLiP focuses on improving the quality of protein–ligand interaction data, as summarised above; while BioLiP2 aims to improve the usability of the database. Therefore, we believe both databases are valuable to the community of protein–ligand interactions.

## Data availability

Q-BioLiP is freely available at: https://yanglab.qd.sdu.edu.cn/Q-BioLiP/.

## CRediT author statement

**Hong Wei**: Data curation, Software, Original draft preparation. **Wenkai Wang:** Original draft preparation. **Zhenling Peng:** Supervision, Original draft preparation. **Jianyi Yang:** Conceptualization, Methodology, Supervision, Writing - Reviewing and Editing.

## Acknowledgments

This work is supported in part by the National Natural Science Foundation of China (Grant Nos: T2225007 and T2222012).

## Conpeting interests

None.

## Supplementary material

**Figure S1.**
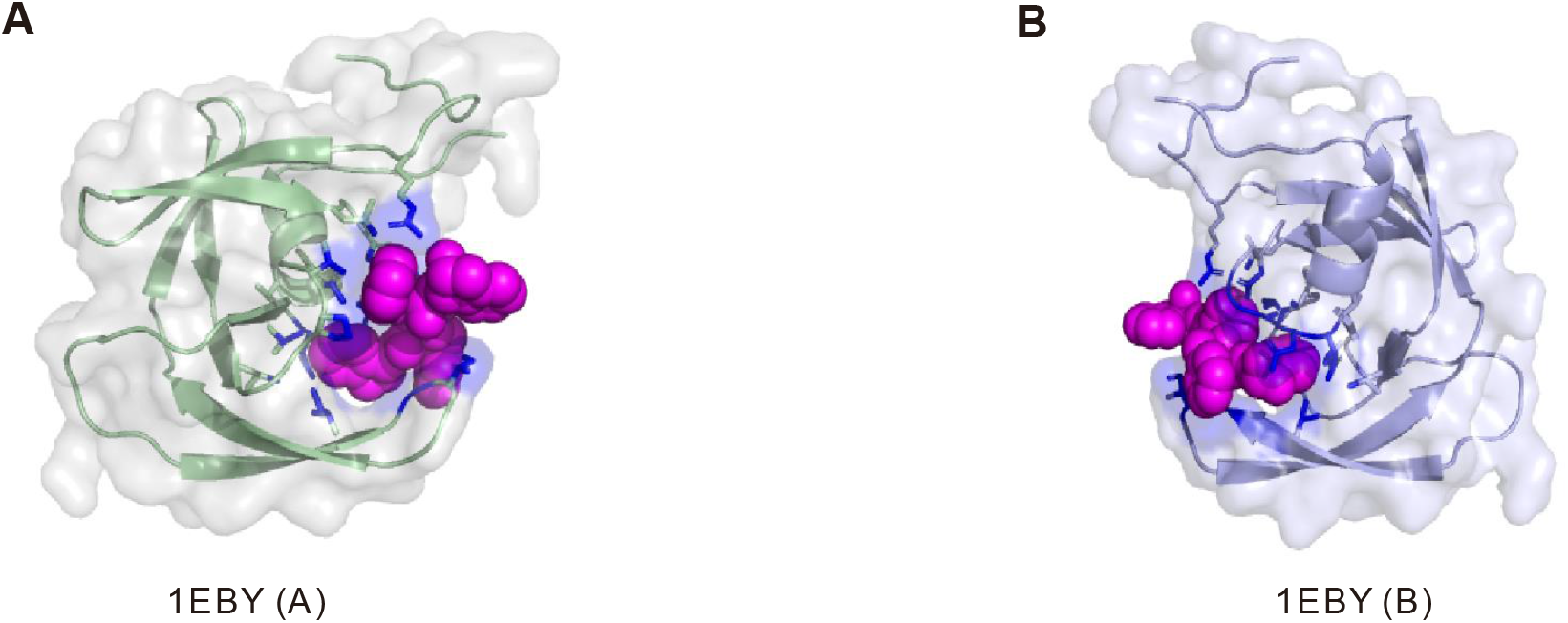
BioLiP entries for an example structure (PDB ID: 1EBY) **A.** The chain A of the HIV-1 protease bound with its inhibitor. **B.** The chain B of the HIV-1 protease bound with its inhibitor.

**Figure S2.**
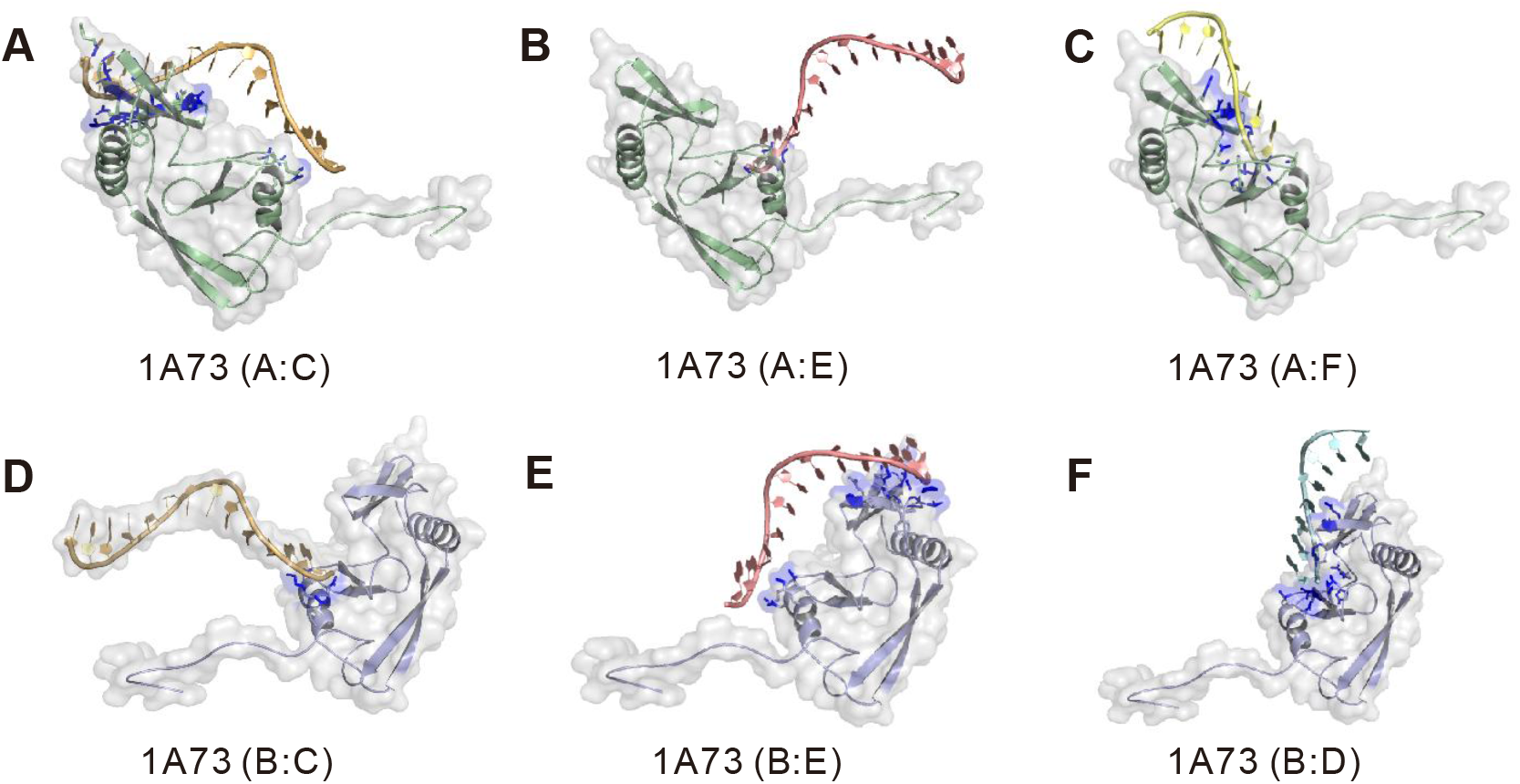
BioLiP entries for an example protein–DNA interaction (PDB ID: 1A73) The DNA chains C (**A**), E (**B**), and F (**C**) bound with the protein chain A, respectively. The DNA chains C (**D**), E (**E**), and D (**F**) bound with the protein chain B, respectively.

**Figure S3.**
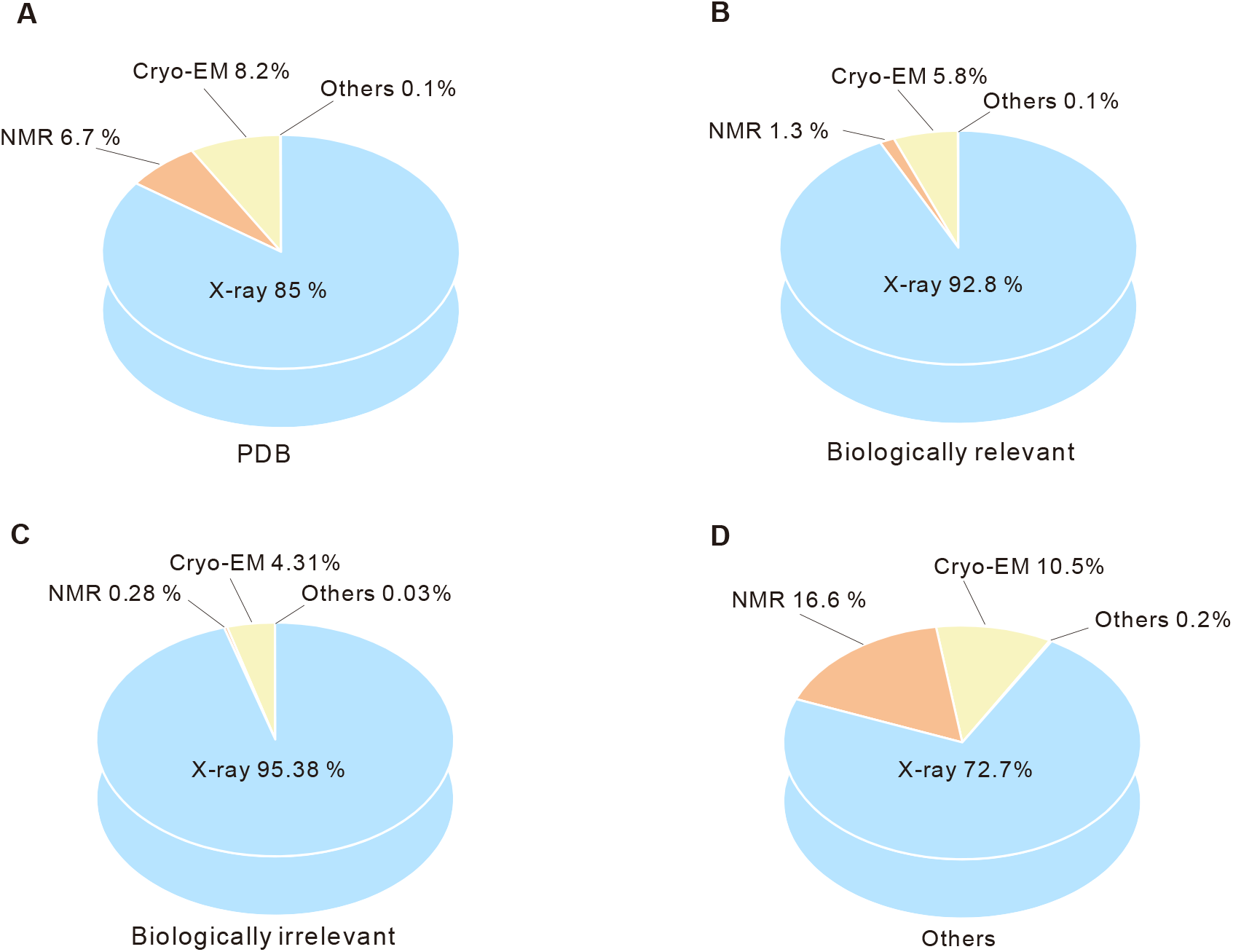
Distribution of structure determination methods. The proportion of structures in PDB (**A**), biologically relevant entries (**B**), biologically irrelevant entries (**C**), and other entries (**D**). NMR, Nuclear Magnetic Resonance; X-ray, X-radiation; Cryo-EM, Cryo-Electron Microscopy. Others includes EPR and neutron diffraction and so on.

**Figure S4.**
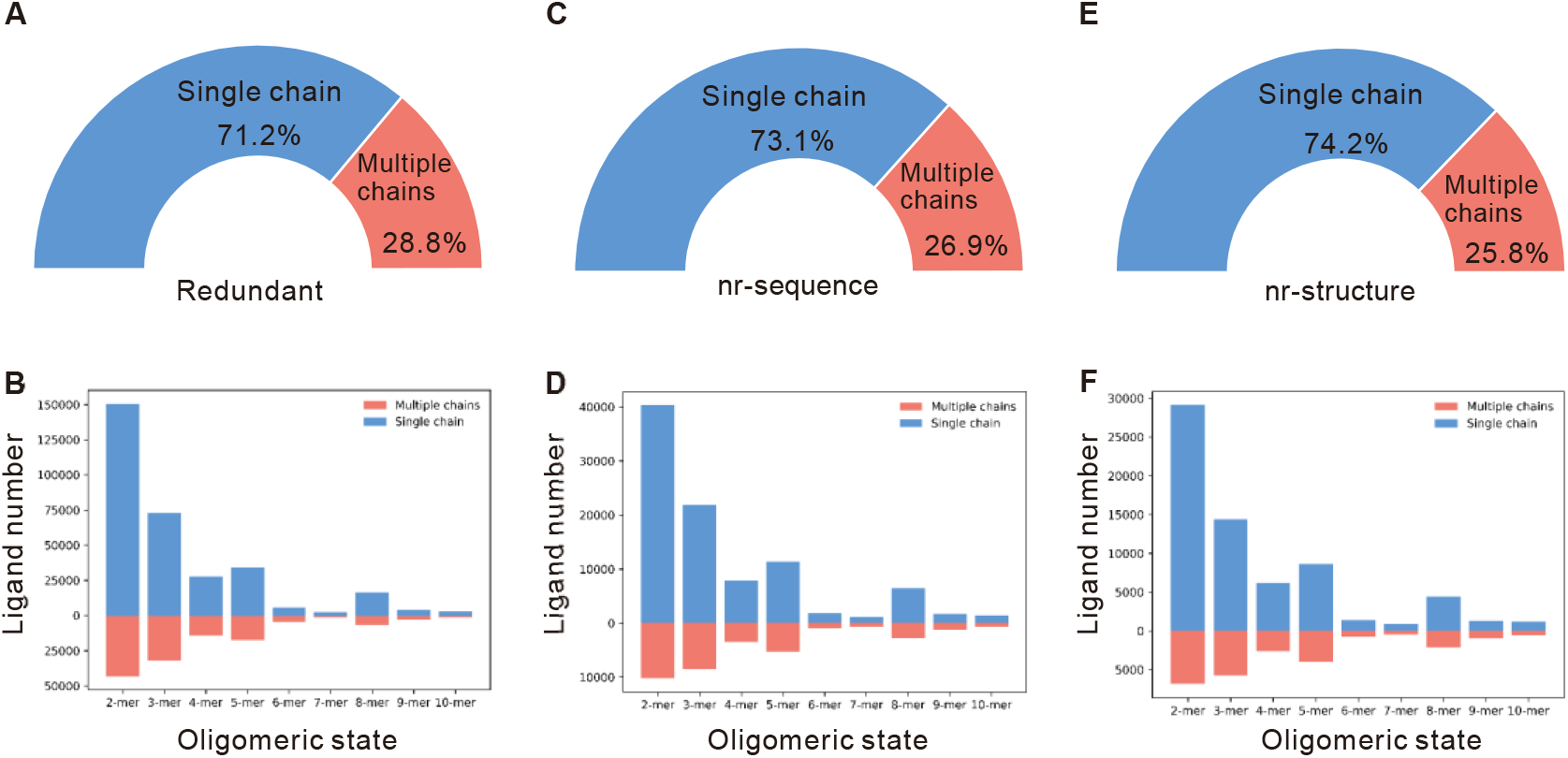
Distributions of the ligand binding data. The proportion of ligand binding with single chain and multiple chains in the redundant (**A**), nr-sequence (**C**), nr-structure datasets (**E**), respectively. The distribution of ligand binding with single chain and multiple chains in redundant (**B**), nr-sequence (**D**), nr-structure datasets (**F**) at different oligomeric states, respectively.

**Figure S5.**
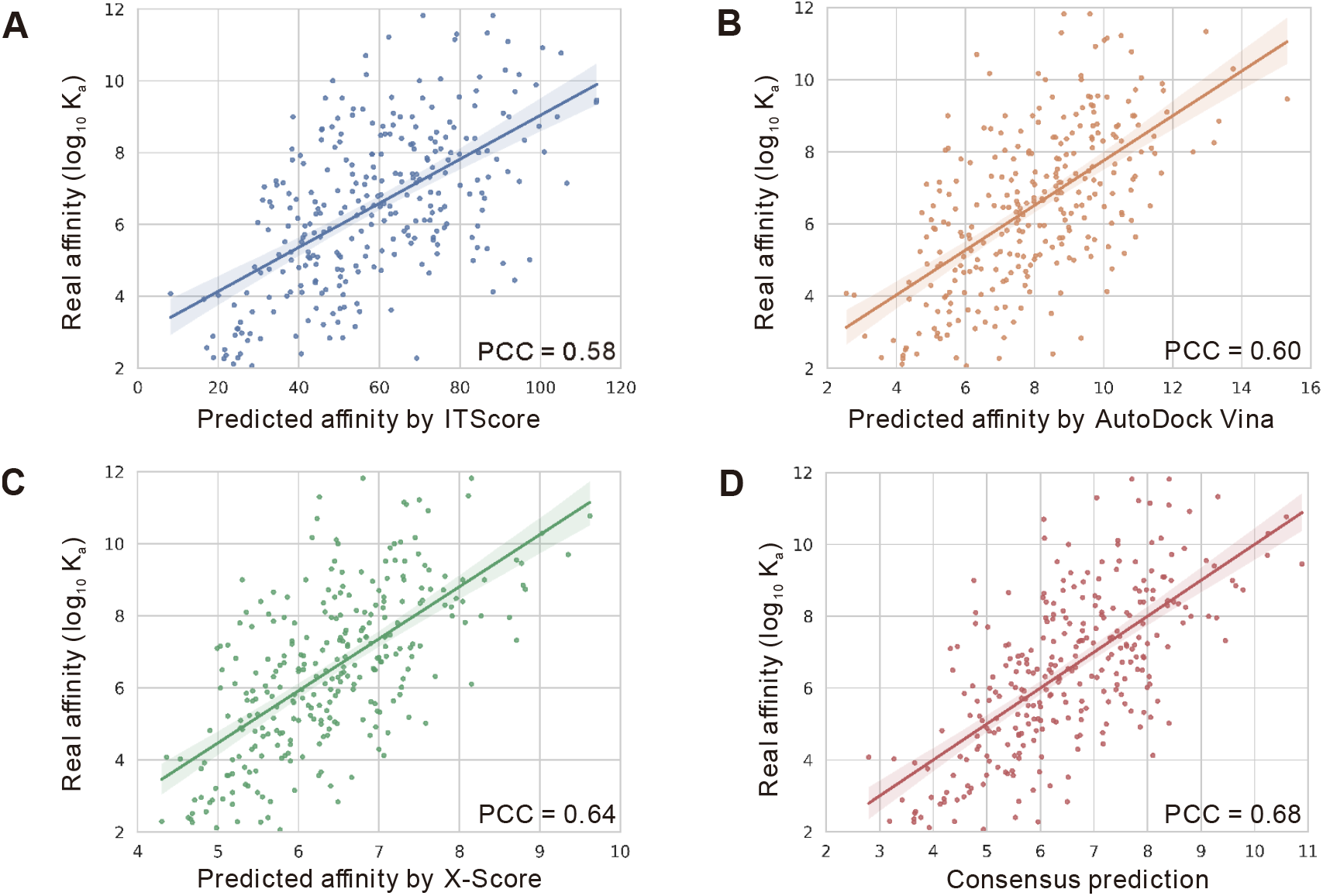
Relationship between the predicted and the real binding affinities. The relationship between the real affinity and ITScore (**A**), AutoDock Vina (**B**), and X-Score (**C**), respectively. **D.** The relation between the real affinity and the consensus prediction.

**Figure S6.**
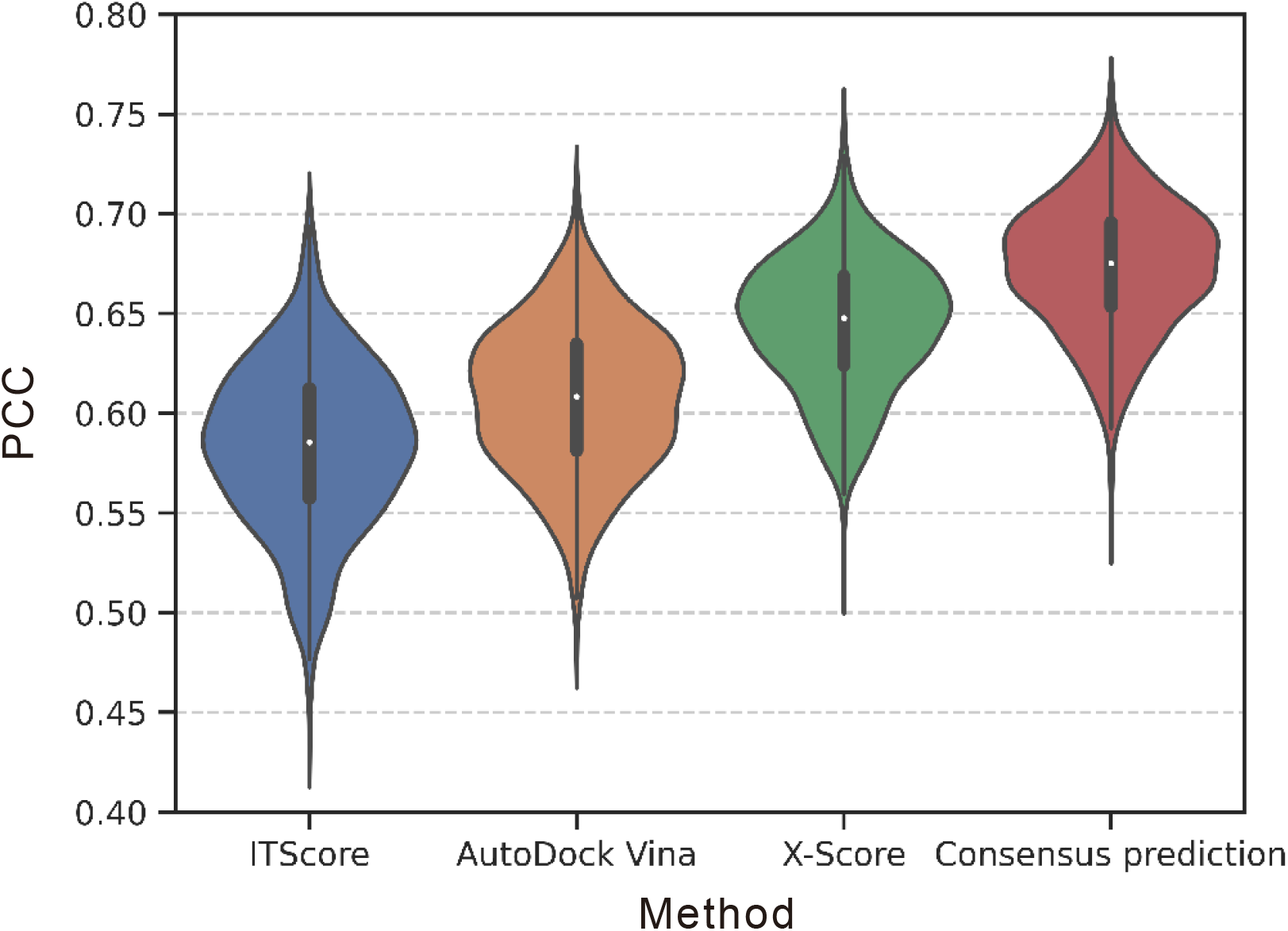
Distributions of the PCCs between predicted and real binding affinities by different methods. The distributions were obtained based on bootstrap sampling (1,000 times).

